# COLSTATS: an interactive web platform for systematic colocalization of GWAS and QTL summary statistics

**DOI:** 10.64898/2026.01.28.702279

**Authors:** Maria Antonietta Diana, Stefano Onano, Marco Masala, Davide Murrau, Francesco Cucca, Federico Santoni, Mauro Pala

## Abstract

Colocalization of GWAS with QTL data is pivotal for identifying shared genetic determinants of molecular traits and diseases yet remains hindered by fragmented data sources and technical barriers. COLSTATS addresses this gap by offering a web-based Shiny platform that integrates 1,648,851 harmonized summary statistic datasets from key resources (GWAS Catalog, UK Biobank, GTEx, BluePrint, eQTLGen, and immune-phenotype studies), all formatted as VCFs aligned to hg38 with uniform chr_pos_ref_alt variant identifiers. Users can interactively select two traits, specify a genomic region or gene, define priors, and execute colocalization via coloc.abf, with results displayed in tabular and graphical form, downloadable as PDF, CSV, or PNG. The system includes result caching, history tracking, and an intuitive interface for streamlined exploration.COLSTATS is freely accessible at https://colstats.irgb.cnr.it. This resource empowers researchers, including non-bioinformaticians, to perform transparent and reproducible colocalization analyses efficiently.

## Introduction

Genome-wide association studies (GWAS) have identified thousands of loci associated with human traits and diseases. A major challenge remains the pinpointing of causal variants, given the extensive linkage disequilibrium across genomic regions and the complexity of traits that may involve multiple interacting factors (1). In parallel molecular-QTL datasets link variants to molecular phenotypes such as gene expression (e.g. eQTLs from GTEx (2), immune-cell QTLs (3), protein abundance (pQTL), methylation (mQTL), splicing (sQTL) and others (4). Colocalization methods integrate GWAS and molecular QTL data to identify variants likely influencing both trait and gene through a shared causal signal, thereby providing mechanistic insights (5). However, harmonizing heterogeneous data sources—each with different formats, genome builds, allele encodings, and metadata—complicates systematic colocalization across the landscape of GWAS/QTL data. Moreover, existing tools often require command-line expertise and bespoke preprocessing, which might limit accessibility for non-computational experts.

We present COLSTATS, an interactive web platform designed to address these challenges. It provides a centralized repository of harmonized, VCF-based GWAS/QTL summary statistics, aligned to hg38, and enables user-friendly, reproducible colocalization analysis in three straightforward steps: selecting traits, specifying a region or gene, and running colocalization using the coloc package. The platform supports flexible priors, provides caching/history features for increased efficiency, and outputs both numerical summaries, local association plots and sensitivity analysis.

## Materials and Methods

### Database data sources and harmonization

For all studies integrated into COLSTATS, we aimed to obtain, whenever available, summary statistics including at least the following fields: chromosome (chr), position (pos), variant identifier, allele information (effect and other allele), effect size (beta), standard error (SE), p-value, allele frequency (AF), and effective sample size (N).) All datasets were harmonized to the GRCh38/hg38 genome build and when summary statistics were provided in other builds, genomic coordinates were converted using LiftOverVcf from GATK v4.1.5.0 (10). The following sections describe the methods applied to each dataset.

*GWAS Catalog – NHGRI-EBI Catalog (PubmedID: 27899670)(6)*. We retrieved GWAS Catalog metadata and summary statistics from http://ftp.ebi.ac.uk. were obtained from the studies metadata file (v1.0.2.1) and from per-study YAML files, which report GWAS-level details including trait description, sample size, and population frequencies. The location of summary statistics was obtained from the harmonised_list.txt file. In total, we downloaded **80**,**295 harmonized summary statistics** files (∼24 Tb, human genome build hg38), each corresponding to a specific trait or disease.

For each GWAS, we retrieved variant-level details (variant name, chromosome, genomic position, effect allele, non-effect allele, and effect allele frequency) and association statistics (beta, standard error, p-value, and sample size). To obtain sample size, we developed a custom script that scans the header of each summary statistics file (converted to lowercase) for a column name matching one of a set of predefined identifiers (e.g., n, samplesize, n_total, neff). If no matching column is found in the summary file, the script searches into the metadata file. We prioritized sample size extraction from the summary statistics because in some studies not all variants were tested in the full set of samples, resulting in variant-specific sample sizes. Extracting sample size directly from the summary statistics allowed us to account for this per-variant variability.

For downstream analysis, we required the following columns to be present: hm_chrom, hm_pos, hm_variant_id, hm_effect_allele, hm_other_allele, hm_beta, standard_error, p_value, and hm_effect_allele_frequency. Overall, we obtained a total of 18,438 **GWAS** summary **statistics** files matching these criteria.

Studies were classified as case–control (cc) or quantitative (quant). Case–control studies were defined as those for which both the number of cases and controls were reported (i.e., not ‘NA’); all others were classified as quantitative.

Summary statistics were harmonized using gwas2vcf (7) (docker image mrcieu/gwas2vcf) with Homo_sapiens.GRCh38.dna.toplevel.fa (from Ensembl) as the reference genome, generating harmonized VCF-formatted results. VCF files were then indexed with tabix (8). Variant identifiers were coded as chr_pos_ref_alt (chromosome, position, reference allele, alternative allele). Finally, we assigned a unique colstats_id to each GWAS by naming the summary statistics with the prefix cs_X_a, where X is an integer identifier.

*UK Biobank - UKBB (PubmedID: 30305743)(9)*. We retrieved UK Biobank metadata by downloading the phenotype flat file from https://pan.ukbb.broadinstitute.org/downloads, which reports GWAS-level details including trait description, sample size, population allele frequencies, and the location of summary statistics. A total of 7,228 GWAS summary statistics files (∼6.7 Tb) were retrieved, each representing a unique trait or disease. Each file included multiple ancestry-specific summary statistics (AFR, AMR, CSA, EAS, EUR, MID), together with a combined meta-analysis across populations (META).

For each GWAS, we extracted variant-level information (variant identifier, chromosome, genomic position, effect allele, non-effect allele, and effect allele frequency) and association statistics (beta, standard error, p-value, and sample size). Variant identifiers were coded as chr_pos_ref_alt (chromosome, position, reference allele, alternative allele). The effect allele was defined as the alternative allele, in accordance with the UK Biobank manifest (https://pan.ukbb.broadinstitute.org/docs/per-phenotype-files/index.html). P-values were reconstructed from neglog10_pval as p = 10^(-neglog10_pval). Studies were classified as either case–control (cc) or quantitative (quant). Case–control studies were defined as those in which both the number of cases and controls were greater than zero; all other studies were classified as quantitative. Sample sizes were extracted from the metadata table. For case–control GWAS, the total sample size was calculated as the sum of cases and controls. For quantitative GWAS, where one of the two fields (cases or controls) was set to zero, the sample size was derived from the non-zero field. Effect allele frequency (EAF) was conservatively estimated. For meta-analysis GWAS with case–control design, we computed a sample-size–weighted average of case and control allele frequencies when available:

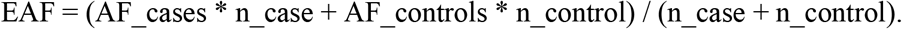

If population-specific allele frequencies were missing, we used the combined meta-level frequency (af_meta or eaf) when provided. For quantitative traits in the meta-analysis, the combined AF was used if available. Summary statistics were harmonized using gwas2vcf (docker image mrcieu/gwas2vcf) with human_g1k_v37.fasta as the reference genome, generating harmonized VCF-formatted GWAS results. Genomic coordinates were then lifted over to hg38 using LiftoverVcf from GATK v4.1.5.0 (10). We retained only standard chromosomes (1–22, X, Y), removed the “chr” prefix, and replaced contig names in the VCF header with Ensembl chromosome notation. VCF files were indexed with tabix. Finally, we assigned a unique colstats_id to each GWAS by naming the summary statistics with the prefix cs_X_b, where X is an integer identifier. Overall, we obtained a total of 12,780 **GWAS summary statistics**.

*Orrù et al. 2020 (PubmedID: 32929287)(3) –* This dataset includes summary statistics of immunophenotypes (traits obtained with citofluorimetric analysis including cell counts, parental percentages and surface protein expression), corresponding to **731 GWAS summary statistics**. Summary statistics of the study were already present in the GWAS Catalog but the alleles of the INDEL were not explicited: they were coded as R/D or R/I. From the SardiNIA imputed panel (11) we retrieved the regular name of the indels alleles which we substituted to the summary statistics. Summary statistics were harmonized with gwas2vcf docker (mrcieu/gwas2vcf) by using human_g1k_v37.fasta as reference genome, which produces a harmonized summary statistic in VCF format. We lifted the coordinated to hg38 by using (LiftoverVcf from gatk-4.1.5.0). We kept only standard chromosomes names (1, .., 22, X and Y) and renamed the chromosome name without “chr” suffix and substituted the contigs names in the header with those of the ensemble chromosome notation. We then indexed the VCF with tabix. For the metadata, the accurate name of the immunotrait tested (e.g. Naive-mature B cell %B cell, IgD+ CD38dim AC, Lymphocyte %leukocyte) was derived from the supplementary table 1B of Orrù et al., 2020. Variant name was coded as chr_pos_ref_alt (chrom, pos, ref and alt stand for chromosome, genomic position, reference allele and alternative allele). To assign a colstats_id, we named the summary statistics of this dataset using the code cs_X_d, where X is an integer number.

**Table 1:**
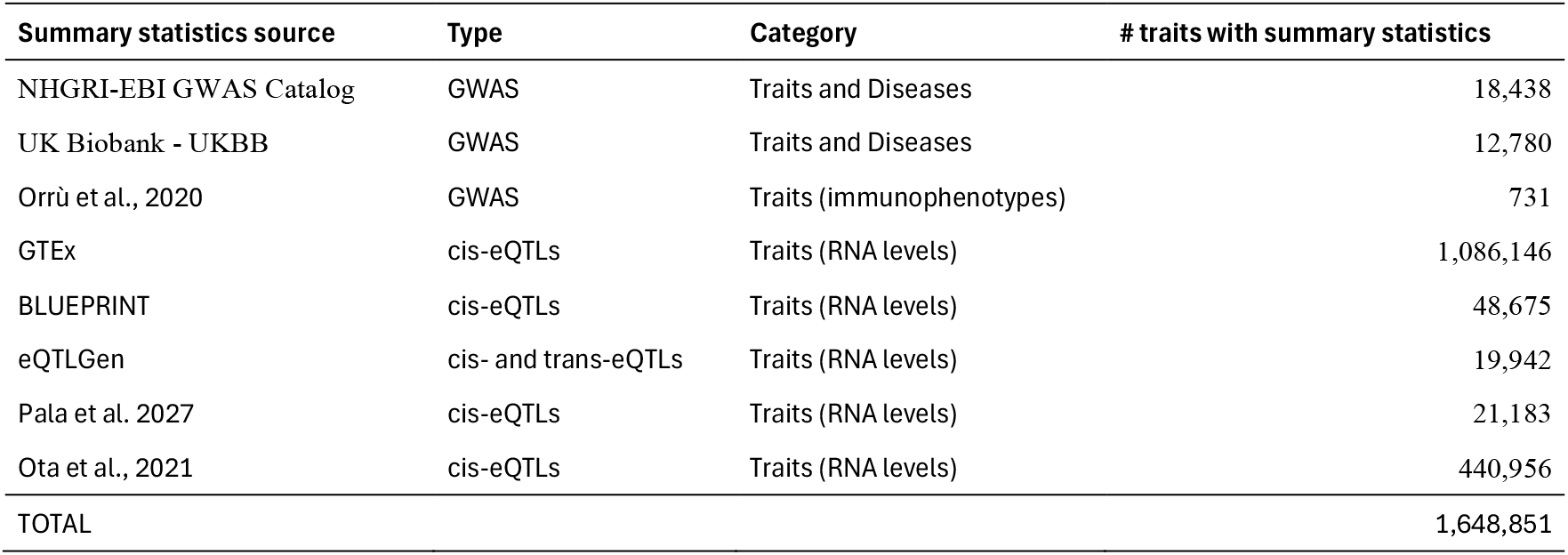
Summary of dataset inventory across sources (e.g., number of summary statistics per resource).

*GTEx eQTLs (PubmedID: 29022597)(2)*We downloaded the summary statistics from (https://gtexportal.org/home/). This dataset includes eQTLs for 44 major human tissues in human genome build 37 including: Adipose Subcutaneous, Adipose Visceral Omentum, Adrenal Gland, Artery Aorta, Artery Coronary, Artery Tibial, Brain Anterior cingulate cortex BA24, Brain Caudate basal ganglia, Brain Cerebellar Hemisphere, Brain Cerebellum, Brain Cortex, Brain Frontal Cortex BA9, Brain Hippocampus, Brain Hypothalamus, Brain Nucleus accumbens basal ganglia, Brain Putamen basal ganglia, Breast Mammary Tissue, Cells EBV-transformed lymphocytes, Cells Transformed fibroblasts, Colon Sigmoid, Colon Transverse, Esophagus Gastroesophageal Junction, Esophagus Mucosa, Esophagus Muscularis, Heart Atrial Appendage, Heart Left Ventricle, Liver, Lung, Muscle Skeletal, Nerve Tibial, Ovary, Pancreas, Pituitary, Prostate, Skin Not Sun Exposed Suprapubic, Skin Sun Exposed Lower leg, Small Intestine Terminal Ileum, Spleen, Stomach, Testis, Thyroid, Uterus, Vagina, Whole Blood. These data corresponded to 1,086,146 cis-eQTLs summary statistics. Summary statistics were harmonized with gwas2vcf docker (mrcieu/gwas2vcf) by using human_g1k_v37.fasta as reference genome, which produces a harmonized summary statistic in VCF format. We lifted the coordinated to hg38 by using (LiftoverVcf from gatk-4.1.5.0). We kept only standard chromosomes (1, .., 22, X and Y) and renamed the chromosome name without “chr” suffix and substituted the contigs names in the header with those of the ensemble chromosome notation. Variant name was coded as chr_pos_ref_alt (chrom, pos, ref and alt stand for chromosome, genomic position, reference allele and alternative allele). To assign a colstats_id, we named the summary statistics of this dataset using the code cs_X_d, where X is an integer number..

*BLUEPRINT eQTLs (PubmedID: 27863251)(12) –*We downloaded the summary statistics from http://blueprint-dev.bioinfo.cnio.es/WP10/qtls. This dataset includes eQTLs for three major human immune cell types (CD14+monocytes, CD16+ neutrophils, and naive CD4+ T cells) from up to 194 individuals (194, 192 and 171 individuals respectively), corresponding to a total of **48**,**675 cis-eQTLs summary statistics**. Summary statistics were harmonized with gwas2vcf docker (mrcieu/gwas2vcf) by using human_g1k_v37.fasta as reference genome, which produces a harmonized summary statistic in VCF format. We lifted the coordinated to hg38 by using (LiftoverVcf from gatk-4.1.5.0). We kept only standard chromosomes (1, .., 22, X and Y) and renamed the chromosome name without “chr” suffix and substituted the contigs names in the header with those of the ensemble chromosome notation. We then indexed the VCF with tabix. Variant name was coded as chr_pos_ref_alt (chrom, pos, ref and alt stand for chromosome, genomic position, reference allele and alternative allele). To assign a colstats_id, we named the summary statistics of this dataset using the code cs_X_d, where X is an integer number.

*eQTLGen phase 1 (PubmedID: 34475573)* (13)–This dataset includes cis- and trans eQTLs from blood-derived expression from 31,684 individuals through the eQTLGen Consortium, corresponding to a total of **39**,**884 summary statistics (19**,**250 for cis and 19**,**942 for trans eQTLs)**. We downloaded the summary statistics from the eQTLgen website (https://eqtlgen.org/phase1.html). Standard Error was not provided in the original summary statistics and was retrieved (following the method suggested in the eQTLgen website https://download.gcc.rug.nl/downloads/eqtlgen/cis-eqtl/SMR_formatted/README_cis_full_SMR) by a python custome script with the following commands

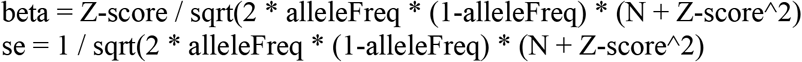

Allelle frequency (alleleFreq) was downloaded from the eQTLgen website (https://download.gcc.rug.nl/downloads/eqtlgen/cis-eqtl/2018-07-18_SNP_AF_for_AlleleB_combined_allele_counts_and_MAF_pos_added.txt.gz)

Summary statistics were then harmonized with gwas2vcf docker (mrcieu/gwas2vcf) by using human_g1k_v37.fasta as reference genome, which produces a harmonized summary statistic in VCF format. We lifted the coordinated to hg38 by using LiftoverVcf from gatk-4.1.5.0. We kept only standard chromosomes (1, .., 22, X and Y) and renamed the chromosome name without “chr” suffix and substituted the contigs names in the header with those of the ensemble chromosome notation. We then indexed the VCF with tabix. Allele frequencies (2018-07-18_SNP_AF_for_AlleleB_combined_allele_counts_and_MAF_pos_added.txt.gz) were downloaded from https://eqtlgen.org/cis-eqtls.html. For each gene, cis- and trans-eQTL summary statistics were combined into a single comprehensive file. Variant name was coded as chr_pos_ref_alt (chrom, pos, ref and alt stand for chromosome, genomic position, reference allele and alternative allele). For the colstats_id, we named the summary statistics of this dataset using the code cs_X_d, where X is an integer number. Overall, we obtained a total of **19**,**942**

### GWAS summary statistics

*Pala et al. 2017 (PubmedID:* 28394350*)(14)* –This dataset includes cis-eQTLs from leukocytes of 606 individuals of the SardiNIA Cohort. We downloaded the summary statistics from https://eqtlsdownload.irgb.cnr.it (full summary statistics with conditional analysis – i.e. primary and after stepwise forward regression) corresponding to a total of **21**,**183 cis-eQTLs summary statistics**. Summary statistics were harmonized with gwas2vcf docker (mrcieu/gwas2vcf) by using human_g1k_v37.fasta as reference genome, which produces a harmonized summary statistic in VCF format. We lifted the coordinated to hg38 by using LiftoverVcf from gatk-4.1.5.0. We kept only standard chromosomes (1, .., 22, X and Y) and renamed the chromosome name without “chr” suffix and substituted the contigs names in the header with those of the ensemble chromosome notation. We then indexed the VCF with tabix. Variant name was coded as chr_pos_ref_alt (chrom, pos, ref and alt stand for chromosome, genomic position, reference allele and alternative allele). To assign a colstats_id, we named the summary statistics of this dataset using the code cs_X_d, where X is an integer number.

*Ota et al*., *2021 (PubmedID: 33930287)(15)*. This dataset consists of 28 distinct immune cell subsets (CD16p Mono, CL Mono, CM CD8, DN B, EM CD8, Fr III T, Fr II eTreg, Fr I nTreg, Int Mono, LDG, Mem CD4, Mem CD8, NC Mono, NK, Naive B, Naive CD4, Naive CD8, Neu, Plasmablast, SM B, TEMRA CD8, Tfh, Th17, Th1, Th2, USM B, mDC, pDC) from 337 patients diagnosed with 10 categories of immune-mediated diseases (systemic lupus erythematosus (SLE), idiopathic inflammatory myopathy (IIM), systemic sclerosis (SSc), mixed connective tissue disease (MCTD), Sjö gren’s syndrome (SjS), rheumatoid arthritis (RA), Behcet’s disease (BD), adult-onset Still’s disease (AOSD), ANCA-associated vasculitis (AAV), or Ta-kayasu arteritis (TAK)) and 79 healthy volunteers. This dataset overall consists of **440**,**956 cis-eQTLs summary statistics**. Summary statistics have been downloaded from NBDC Human Database (https://ddbj.nig.ac.jp). As standard errors were not available in the released summary statistics, they were inferred from the reported effect sizes, p-values, and sample sizes for each cell type. With a custom R script with the following commands:

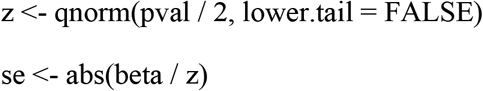

Sample size information was retrieved from the Ota_2021 web browser (https://www.immunexut.org).

Summary statistics were then harmonized with gwas2vcf docker (mrcieu/gwas2vcf) by using human_g1k_v37.fasta as reference genome, which produces a harmonized summary statistic in VCF format Variant name was coded as chr_pos_ref_alt (chrom, pos, ref and alt stand for chromosome, genomic position, reference allele and alternative allele). To assign a colstats_id, we named the summary statistics of this dataset using the code cs_X_d, where X is an integer number.

### Web Application Architecture

The COLSTATS platform was implemented as an R/Shiny application with a modular client–server architecture.

The frontend was designed with `shinythemes, shinyjs`, custom CSS, and JavaScript to guide users through a clear three-step workflow. Each step is displayed in a dedicated “step card” interface with large navigation tabs and contextual help icons.

In Steps 1 and 2, users specify traits or diseases of interest via a searchable `selectizeInput` field. Upon entry, a metadata table (rendered with `DT` and the *Buttons* extension) summarizes key information for each study, including identifiers (COLSTATS ID, original study ID, PubMed ID), population ethnicity, consortium, study design (case– control or quantitative), and sample size (with separate numbers for cases and controls when applicable). This metadata-driven interface enables users to refine their selection and filter studies according to the most relevant criteria. A summary panel then displays the two selected traits side by side.

In Step 3, users define the input parameters for colocalization. The genomic region can be entered manually or selected by gene (with boundaries from GENCODE v41), with interactive zoom controls (±100 kb) and automatic validation of format and size (maximum 5 Mb). Prior probabilities are entered in dedicated fields with inline help. Once the analysis is launched, progress is tracked by a loading overlay and status indicators. The results include:

i. a *Selection Overview* summarizing traits and genomic region,
ii. colocalization summary tables reporting posterior probabilities (PP.H0–H4), top associated variants with effect sizes and p-values, (iii) sensitivity analyses displayed as plots.

Additional features include an *Analysis History* panel showing the last 10 analyses (with trait names, region, PP.H4, timestamp) from which users can restore results, and multiple download options: a full PDF report (including parameters, tables, and plots), a summary CSV file, and plots exported in PNG or PDF format; and a **“Use example” buttons** that pre-populates the interface with the demonstrative case study described below (RA GWAS vs. CD40 cis-eQTL), allowing new users to immediately explore a complete colocalization workflow without manual trait selection.

On the server side, the user-selected datasets are passed to the `coloc.abf` function to compute posterior probabilities for five mutually exclusive hypotheses (H0–H4). The software automatically determines from metadata whether the trait is quantitative (quant) or case–control (cc). To optimize performance, results are cached and reused if the same inputs are selected. Robust error handling captures invalid genomic regions, missing VCF files, or harmonization inconsistencies, with descriptive feedback returned to the user.

The application integrates core R packages including: `coloc` (for colocalization inference), `VariantAnnotation` and `GenomicRanges` (for parsing and subsetting VCF files), `gwasglue` (for harmonization), `shinyjs` (for interactivity), `DT` (for dynamic tables), and `ragg` (for device-independent graphics rendering).

## Results

### Database Coverage

The COLSTATS platform integrates a comprehensive collection of GWAS and QTL datasets into a unified, harmonized resource. The full database comprises 1,648,851 summary statistics, harmonized with the gwas2vcf docker (mrcieu/gwas2vcf) (7) and indexed with tabix (8). Variants are consistently aligned across studies to ensure uniform allele orientation and identifier encoding. The metadata table provides detailed descriptors, including COLSTATS ID, original study ID, PubMed ID, trait or disease name, sample size (with case/control numbers when applicable), population, year of publication, consortium, author, and study type (quantitative or case– control).

### User Interface and Workflow

Users may interact with COLSTATS via a streamlined three-step process: (i) select trait 1 and trait 2, (ii) specify a genomic region or gene of interest, (iii) adjust prior probabilities and execute colocalization. Results are returned instantly in interactive tables and plots.

### Performance

With tabix indexing, typical colocalization analyses complete in seconds, depending on region size and dataset selection.

### Demonstrative Case Study

To illustrate COLSTATS workflow, we analyzed the CD40 locus by pairing rheumatoid arthritis (RA) GWAS summary statistics from the GWAS Catalog (GCST000679) as Trait 1 with CD40 cis-eQTLs in leukocytes from Pala et al. (N=606) as Trait 2 (Figure 2 and 3). This analysis is also available through the “Use example” buttons in the web interface, enabling users to reproduce these results interactively. Within the Pala resource, two eQTLs are available for CD40: an unconditioned scan (Conditional0) and a conditionally independent signal obtained by forward stepwise regression (Conditional1). We selected Conditional1 as Trait 2. (Figure 3). After defining the CD40 locus as the genomic window of interest, colocalization was performed with **coloc.abf** under default priors (Figure 4).

**Figure 1.**
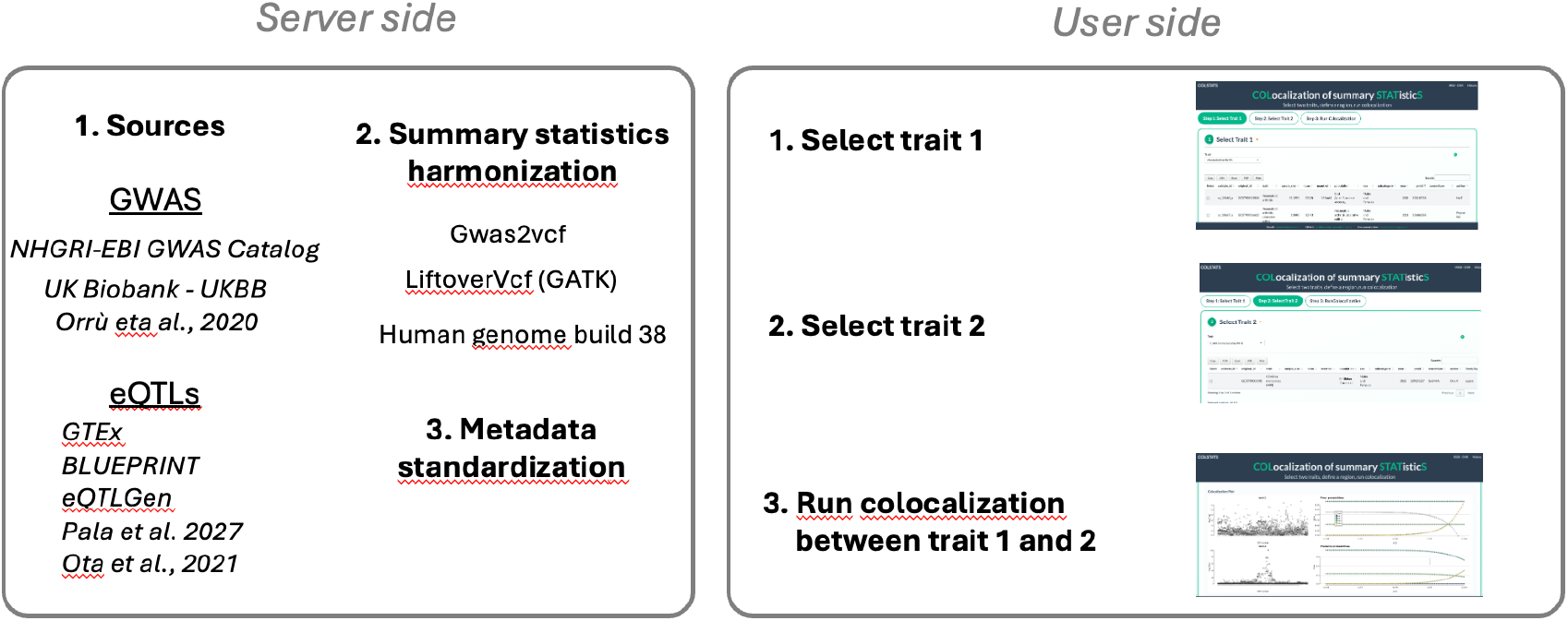
Server-side and user-side workflow of COLSTAT. On the **server side**, summary statistics from multiple sources (GWAS Catalog, UK Biobank, population cohorts, and eQTL datasets such as GTEx, BLUEPRINT, and eQTLGen) are collected. These data are harmonized through standardized variant representation (e.g., gwas2vcf, liftover to the human reference genome build 38) and metadata unification. On the **user side**, the web interface guides the analysis in three steps: (1) selection of the first trait of interest, (2) selection of the second trait, and (3) colocalization analysis between the two traits. The results include interactive regional plots and posterior probability estimates that support causal inference.

**Figure 2.**
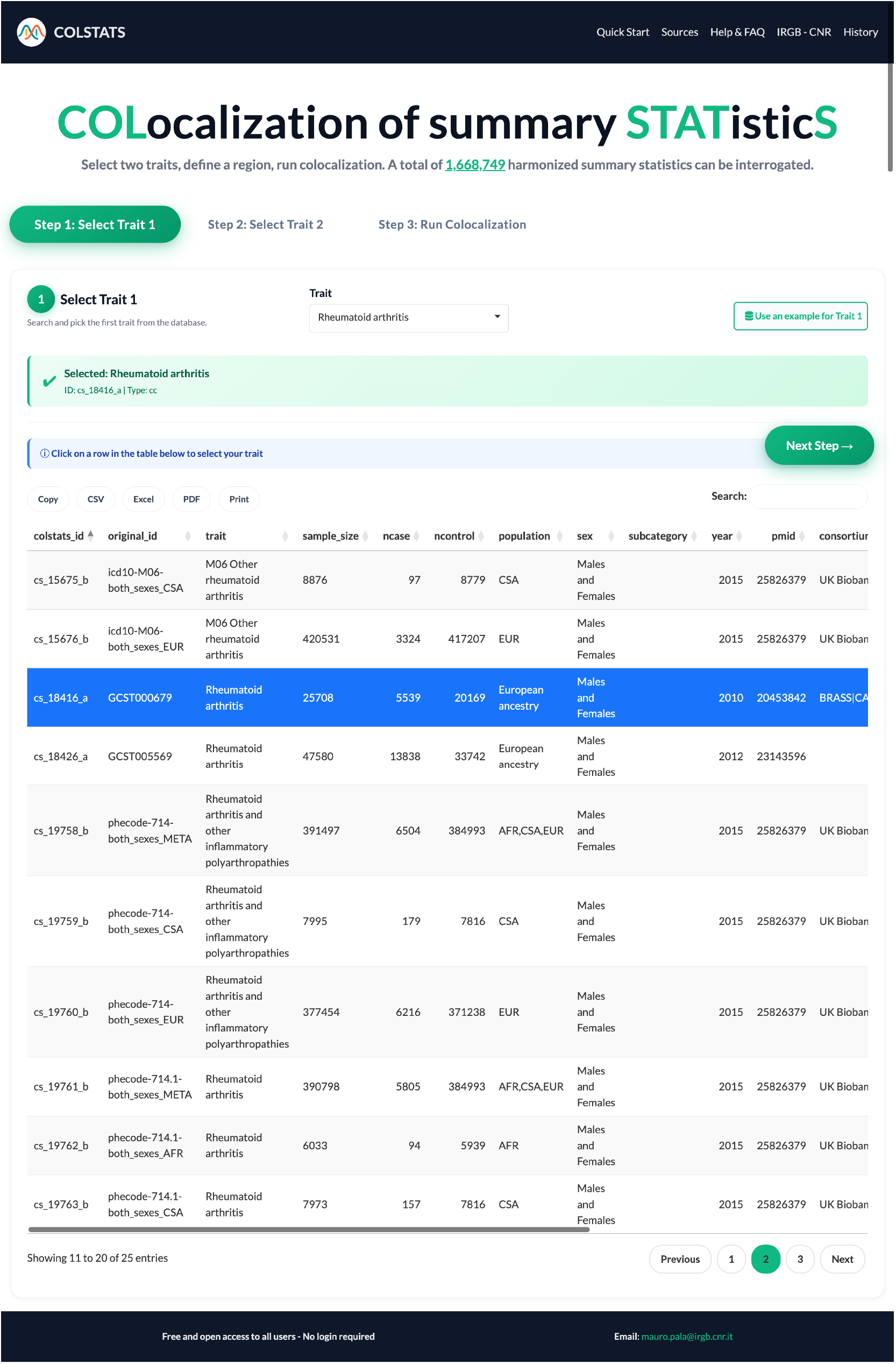
Step 1—Select Trait 1 in COLSTATS. The interface displays a searchable, sortable table of curated GWAS summary-statistics records from which the user chooses the disease/trait to be used in the colocalization workflow. Columns shown include colstats_id, original_id, trait, sample_size, ncase, ncontrol, population, sex, subcategory, year, PMID, and consortium, with one-click export options (CSV/Excel/PDF). In this example, rheumatoid arthritis is selected (blue highlight; GCST000679), which will be carried forward to Steps 2–3 to test colocalization with the CD40 cis-eQTL measured in peripheral blood leukocytes.

**Figure 3.**
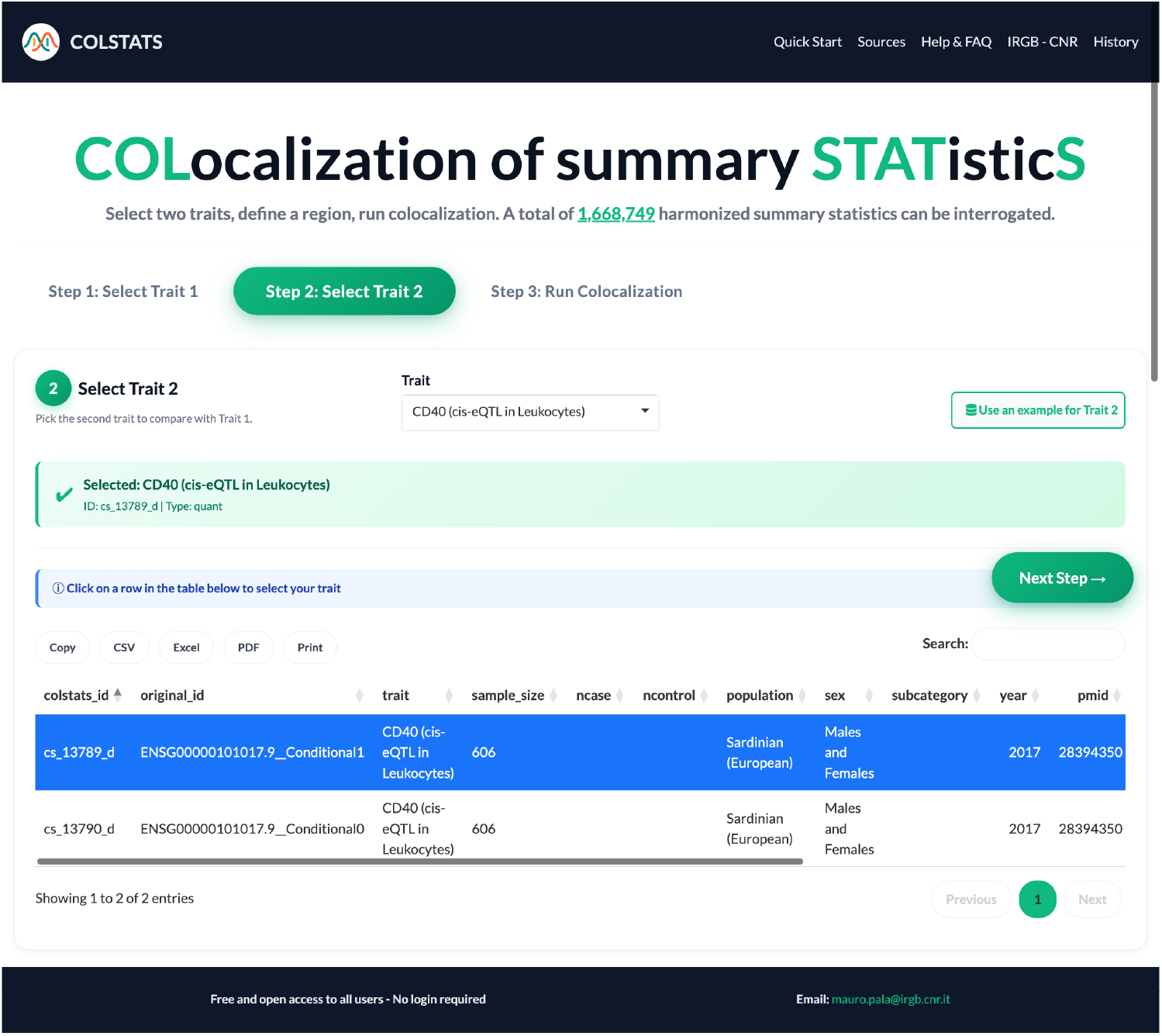
Step 2—Select Trait 2 in COLSTAT . The interface lists molecular traits available for colocalization. Here, CD40 cis-eQTL in leukocytes from Pala et al (14) was chosen. Within the Pala leukocyte eQTL resource, two versions are currently available for CD40: (i) the unconditioned scan (*Conditional0*; no conditional analysis) and (ii) a conditionally independent signal (*Conditional1*) obtained by forward stepwise regression. In this example, the conditional dataset is selected (blue highlight; *colstats_id* cs_734_d, *original_id* ENSG00000101017.9 Conditional1). Metadata shown include sample size (N=606), population (Sardinian/European), sex (males and females), year (2017), PMID (28394350), and consortium (SardiNIA). Other eQTL resources may be added to COLSTAT in future releases; this figure illustrates the options currently available for CD40 in the Pala dataset and defines Trait 2 for downstream colocalization with the rheumatoid arthritis GWAS chosen in Step 1.

**Figure 4.**
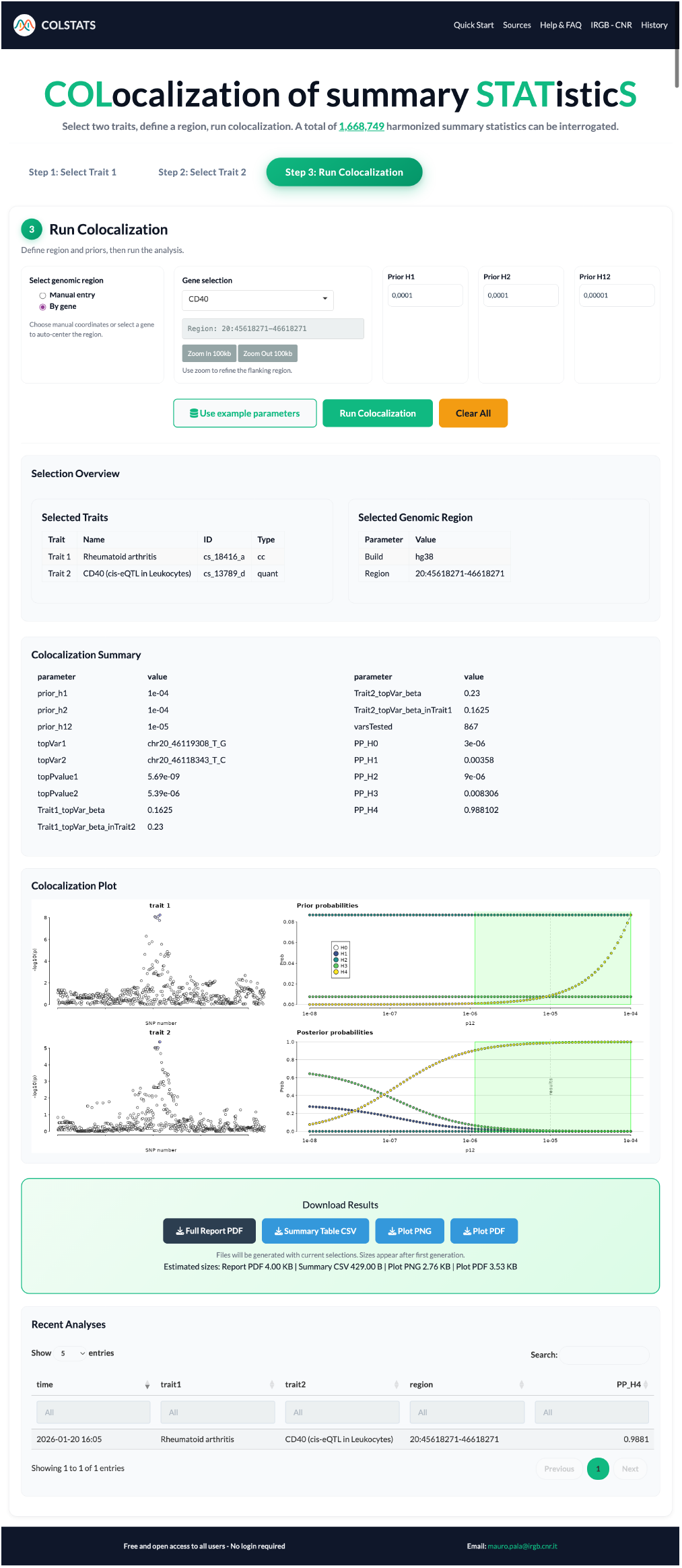
Step 3—Run Colocalization in COLSTATS and visualize results. The CD40 locus is selected because we test a cis-eQTL effect on CD40 for colocalization with rheumatoid arthritis (RA) GWAS (Trait 1). Default priors were used (prior H1 = 1×10^−4^, prior H2 = 1×10^−4^, prior H12 = 1×10^−5^), and 867 variants were evaluated in the region. The strongest RA association is at chr20:46119308_T_G (Trait 1 top P = 5.69×10^−9^), while the strongest CD40 eQTL is at chr20:46118843_T_C (Trait 2 top P = 5.39×10^−6^). Colocalization results (coloc): Posterior probabilities are PP_0_=3×10^−6^, PP_1_=0.00358, PP_2_=9×10^−6^, PP_3_=0.008306, and PP_4_=0.988102, providing strong evidence for one shared causal variant driving both the RA signal and the CD40 cis-eQTL. Effect directions for the lead variants are concordant across traits: the RA top variant has a positive effect on CD40expression (β in Trait 2 ≈ +0.23), and the eQTL top variant shows a positive effect on RA (β in Trait 1 ≈ +0.16). This sign concordance indicates that higher CD40 expression associates with increased RA risk. Plots (from the coloc tool): Bottom-left, local association (-log_10_P) plots for Trait 1 (top panel) and Trait 2 (bottom panel) show peaks aligned in the same LD block, visually supporting colocalization. Bottom-right, the sensitivity analysis shows prior (top) and posterior (bottom) probabilities across a range of shared-causal priors (p_12_); PP_4_ remains the dominant outcome over a broad, plausible prior range (vertical line marks the chosen prior; shaded area highlights recommended values), indicating that the inference of colocalization is robust to prior choices.

The strongest RA association in the region was at chr20:46119308_T_G (P = 5.69×10^−9^), and the top CD40 eQTL was chr20:46118843_T_C (P = 5.39×10^−6^). The analysis yielded PP.H4 = 0.988 (PP_4_), providing strong evidence for a shared causal variant between RA and CD40 expression. Cross-trait effect directions for the lead signals were concordant: the RA lead variant increased CD40 expression (Trait-2 β ≈ +0.23), and the eQTL lead variant showed a positive effect on RA (Trait-1 β ≈ +0.16). This sign concordance indicates that higher CD40 expression is associated with increased RA risk as previously observed (17). Results are presented in COLSTATS both tabularly (posterior probabilities; top variants, effect sizes (betas) and P-values) and graphically. Local association plots (from coloc) display aligned −log_10_P peaks across traits within the same LD block, visually supporting colocalization. Moreover, sensitivity analysis shows that PP_4_ remains dominant across a broad range of shared-causal priors, indicating robust inference.

## Discussion

COLSTATS supports researchers bridge the gap between data availability and accessibility by unifying disparate GWAS and QTL resources into a harmonized, user-friendly platform for systematic colocalization analysis. By providing uniformly processed, hg38-aligned VCF summary statistics with consistent variant identifiers, COLSTATS removes a major barrier to integrative analyses.

Compared to traditional approaches such as command-line use of the coloc R package or bespoke pipelines, COLSTATS offers an accessible, zero-install web interface. This broadens access to colocalization analyses, enabling both computational and experimental researchers to explore cross-trait genetic mechanisms.

A unique feature of COLSTATS is the reporting of effect sizes for top variants across traits. This enables users to qualitatively assess whether traits share effects in the same direction (e.g., increased Trait 1 associates with increased Trait 2) or in opposite directions (e.g., increased Trait 1 associates with decreased Trait 2). While formal inference on causal relationships between traits is better addressed by complementary approaches such as Mendelian Randomization (16), the directional consistency of top local variants provides valuable first-line hypotheses regarding the role of genetic and regulatory elements in the locus.

In summary, COLSTATS combines comprehensive data integration, harmonization, and interactive visualization in a single platform, enhancing transparency, reproducibility, and accessibility of colocalization analyses at scale.

## Data Availability

COLSTATS platform: https://colstats.irgb.cnr.it

## Acknowledgments

Funding from Programma di ricerca CN00000013, National Centre for HPC, Big Data and Quantum Computing of the Next Generation EU initiative

